# Vast diversity of anti-CRISPR proteins predicted with a machine-learning approach

**DOI:** 10.1101/2020.01.23.916767

**Authors:** Ayal B. Gussow, Sergey A. Shmakov, Kira S. Makarova, Yuri I. Wolf, Joseph Bondy-Denomy, Eugene V. Koonin

## Abstract

Bacteria and archaea evolve under constant pressure from numerous, diverse viruses and thus have evolved multiple defense systems. The CRISPR-Cas are adaptive immunity systems that have been harnessed for the development of the new generation of genome editing and engineering tools. In the incessant host-parasite arms race, viruses evolved multiple anti-defense mechanisms including numerous, diverse anti-CRISPR proteins (Acrs) that can inhibit CRISPR-Cas and therefore have enormous potential for application as modulators of genome editing tools. Most Acrs are small, highly variable proteins which makes their prediction a formidable task. We developed a machine learning approach for comprehensive Acr prediction. The model showed high predictive power when tested against an unseen test set that included several families of recently discovered Acrs and was employed to predict 2,500 novel candidate Acr families. An examination of the top candidates confirms that they possess typical Acr features. One of the top candidates was independently tested and found to possess anti-CRISPR activity (AcrIIA12). We provide a web resource (http://acrcatalog.pythonanywhere.com/) to access the predicted Acrs sequences and annotation. The results of this analysis expand the repertoire of predicted Acrs almost by two orders of magnitude and provide a rich resource for experimental Acr discovery.

## Introduction

All life forms evolve under constant pressure from numerous, diverse viruses and other parasitic genetic elements and thus have evolved multiple defense systems^1^. The CRISPR-Cas are adaptive immunity systems that are present in nearly all archaea and about 40% of bacteria, and have been harnessed for the development of the new generation of genome editing and engineering tools^2–4^. In the incessant host-parasite arms race, viruses evolved multiple anti-defense mechanisms including numerous, diverse anti-CRISPR proteins (Acrs) that are currently known to comprise 46 distinct families^5, 6^. The Acrs employ different mechanisms to abrogate the activity of CRISPR-Cas systems^7–10^. The majority of the Acrs that have been studied in detail to date bind to functionally important sites of CRISPR-Cas effector proteins and display high specificity towards a particular CRISPR-Cas variant from a narrow range of bacteria or archaea. Some Acrs, however, have broader specificity^11^, for example, acting as nucleic acid mimics^12^. Furthermore, recently, enzymatically active Acrs, such as acetyltransferases and nucleases, have been discovered^13–15^. Clearly, Acrs have enormous potential for application as modulators of genome editing tools^16, 17^. Despite the fundamental interest of Acrs for understanding the biology of host-parasite interactions in prokaryotes and their potential to transform the use of CRISPR in DNA editing, the discovery of novel Acrs remains a formidable task. The principal causes of these difficulties are the small size and extreme evolutionary variability of most of the Acrs which hampers their detection with even the most powerful sequence analysis methods^10^. The currently known Acr families have been discovered using a variety of creative approaches, the two primary ones being guilt-by-association and self-targeting^5, 12, 18–20^.

Guilt-by-association involves searching for homologs of HTH-containing proteins that are typically encoded downstream of previously discovered Acrs^18^. Such proteins are known as anti-CRISPR associated (Aca) and are notably more conserved among viruses than Acrs themselves which greatly facilitates their detection. The genomic neighborhoods encoding Aca homologs are then searched for potential novel Acrs.

Self-targeting genomes are prokaryotic genomes that encode functional CRISPR-Cas systems that encompass spacers targeting regions of their own genome^20^. In this case, CRISPR-Cas system should, in theory, target and kill the host cell. Thus, organisms with self-targeting genomes can only survive when they also carry Acrs that prevent CRISPR-Cas from functioning and keep the cell viable.

Despite the notable success of these two approaches, buttressed by experimental validation of the predictions, neither provides a systematic methodology to detect novel Acrs. The main challenge in discovering novel Acrs is that, in addition to their extreme sequence variability, they share few distinguishing characteristics or similarities outside of their common role in thwarting CRISPR. With no clear way to discern Acrs from other proteins, and no sequence similarity between different Acr families, discovering novel Acrs remains a formidable challenge^21^.

Here, we describe a systematic machine learning approach we developed to predict novel Acrs, based on the few known Acr attributes and a secondary screen using heuristics of known Acrs, to further enrich for likely Acr candidates. We show that this method is significantly predictive of novel Acrs, compile a collection of 2,500 previously undetected predicted Acrs families and examine in detail the top candidates.

## Results

### Characteristic features of the known Acrs

The general concept behind our approach is that we strive to combine the few characteristics Acrs tend to share into a detection model. Our first step was therefore to assemble and quantify features that previously discovered Acrs appear to have in common. To keep track of the known Acrs, we relied on a combination of curated Acr databases^22, 23^, and our own manual data curation (Supplementary Table 1). At the time of our data curation, 39 Acr families were known (Supplementary Table 1). We used this original set to iteratively search for homologs in the non-redundant (NR) database at the NCBI using PSI-BLAST (see Methods for details) and to construct a multiple protein sequence alignment for each Acr family.

Each of these alignments was then PSI-BLASTed against our local dataset^24^ that includes prokaryotic and prokaryotic virus proteins and consists of a total of 182,561,570 proteins (see Methods for details). All hits with an e-value below the threshold of 10e-4 were manually curated to eliminate obvious false positives, such as partial hits to very large proteins or hits to proteins with unambiguously assigned functions. The final positive set consisted of 3,654 Acrs, spanning 32 families (7 of the known Acr families were not represented in our database; Supplementary Table 1; Supplementary File 1).

The most striking common feature of the Acrs is their small size (weighted mean Acr length: 104aa, Table 1) and the tendency to form sets of small proteins that are encoded by co-directional and closely spaced genes in (pro)virus genomes (hereafter directons) (Figure 1, Table 1). We hypothesize that these directons are largely made up of co-transcribed early anti-defense genes. Acrs are also typically encoded upstream of proteins containing an HTH domain, the Acas^18^.

**Figure 1.**
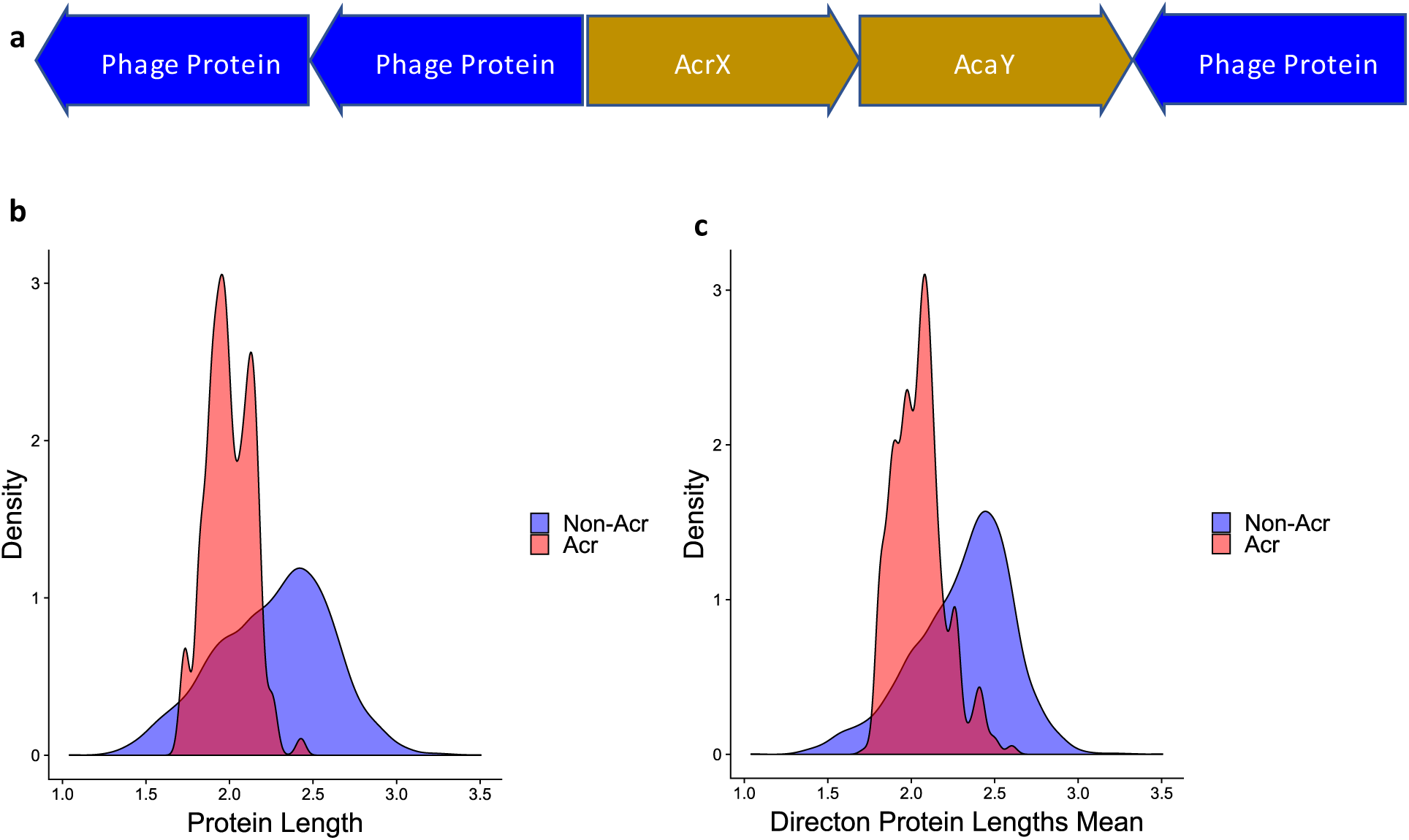
Characteristics of known Acrs. A) A cartoon of a sample directon. Acr proteins characteristically fall upstream of an HTH-domain containing gene, termed Aca. Acrs are usually found in suspected mobile-genetic elements, such as phages. The Acr directon is highlighted in the gold color, while the surrounding proteins are indicated in blue. Characteristically, Acrs fall in directons with small, unidentified proteins. B) A density plot of Acr lengths. C) A density plot of Acr directon mean lengths.

**Table 1.** Feature set. Weighted means of all assessed features, and whether they were used in the final model.

| Feature Name | Acr Mean | Non-Acr Mean | Used in final model |
| --- | --- | --- | --- |
| Containing Genome is Prokaryote | 0.895643762 | 0.895769231 | No |
| Containing Genome is Self-Targeting | 0.334446946 | 0.191923077 | Yes |
| Directon Annotated Protein Fraction | 0.22 | 0.69 | Yes |
| Directon Protein Lengths Mean | 119.27 | 251.71 | Yes |
| Directon Predicted Membrane-Associated Fraction | 0.06 | 0.26 | No |
| Directon Size | 3.49 | 3.5 | Yes |
| Protein is Annotated | 0.064102564 | 0.673076923 | Yes |
| Protein has HTH-Downstream | 0.400757737 | 0.118076923 | Yes |
| Protein is Predicted Membrane-Associated (TMHMM, SignalP) | 0.025641026 | 0.278076923 | No |
| Directon Spacing | 18.37 | 13.7 | No |
| Protein Length | 104.11 | 245.54 | Yes |
| Mean Hydrophobicity (Kyte and Doolittle) | -0.48 | -0.15 | Yes |

Beyond these distinctive features, we considered other protein characteristics that we suspected might be predictive, such as the protein spacing within a directon (Directon Spacing, Table 1) or protein hydrophobicity^25^ (Mean Hydrophobicity, Table 1). We also considered whether proteins gave any significant hits to conserved domains from either CDD^26^ or pVOG^27^ (Is Annotated, Table 1). In total, we constructed a set of 12 features (Table 1, see Methods for details) that, together, provided a compendium of quantifiable features that were used to identify Acr candidates.

### Training and test sets

To build a predictive model, a training set comprised of two components was required: a positive set, consisting of previously discovered Acrs, and a negative set, consisting of proteins confidently inferred not to be Acrs (non-Acrs). For the positive set, the Acrs were weighted by their family and interfamily similarities (Supplementary File 1, Methods), to ensure that related and highly similar Acrs were not over-represented in the training dataset.

Because there is no well-defined, standard set of known non-Acr proteins, we constructed the negative set by randomly selecting viral and prokaryotic proteins, under the assumption that the majority of proteins are non-Acrs. The negative training dataset was constructed by randomly selecting proteins from a combination of 1000 randomly selected prokaryotic virus genomes and 4000 randomly selected CRISPR-Cas-containing prokaryote genomes. Similar to the positive set, we sought to avoid oversampling particular protein families. Therefore, these proteins were clustered by sequence similarity, and for each cluster, a single representative was selected (see Methods for details). We randomly selected 3,500 proteins from this set to constitute the negative, non-Acr set.

During our work on the model, an additional set of Acrs has been published^28, 29^. We incorporated these into our analysis as an unseen test set, i.e. a set of Acrs unavailable during the training stage that we could use to test our model against. Thus, our training set consisted of all known Acrs published before September 2018 (Supplementary File 1; positive set: n=2,775, 26 families; negative set: n=2,600), and the test set consisted of the Acrs published after that date (Supplementary File 1; positive set: n=879 proteins, 6 families; negative set: n=600 proteins).

### Building and evaluating a predictive model

Given our relatively small positive set, we sought to identify a model that would tend towards low variance. To this end, we chose a random forest of extremely randomized trees^30^. As an ensemble method with a highly random component, it has less variance and is therefore less likely to overfit.

The model consisted of a random forest with 1000 decision trees. When training the model, each decision tree is built based on a random sampling of the training data. Each split in the decision tree is determined by randomly selecting multiple values across a random subset of the features, and then setting the values that minimize Gini impurity as the thresholds for the decision tree split. Thus, the final forest consists of 1000 decision trees, where each decision tree’s leaf nodes correspond to members of the training set.

When using the model to assess a candidate protein, the candidate traverses each decision tree. Within each tree, it ends up in a leaf node that contains some mixture of Acrs and non-Acrs from the training set. The tree assigns the candidate a score that is equal to the fraction of Acrs in its leaf (this can be any value between zero and one, inclusive). The score assigned by the model is the mean of the scores across all 1000 trees.

Using the model and the training set we developed, we assessed the performance of the model by 5 iterations of 3-fold cross-validation. In each iteration, the model was trained on two-thirds of the Acr families, and capacity to predict the families that were left out was assessed. For each protein in the test set, we predicted the likelihood of a protein being an Acr using our random forest model. Given the large class imbalance, we down-weighted the negative set in training the model, so that its combined weight was equal to that of the positive set. This weighting was applied to both model training and assessment.

We relied on receiver operating characteristic (ROC) area under the curve (AUC) to assess the model performance and used a genetic algorithm for feature selection. On average, across all 15 cross-validation iterations, we found that our method was significantly predictive of Acrs with an AUC of 0.93 (permutation p-value: 0.001, see Methods for details).

We next used the model to predict Acrs in the unseen test set. The model was found to significantly distinguish Acrs from non-Acrs, with an AUC of 0.83 (permutation p-value: 0.001; Figure 2). This result indicates that our method is indeed predictive of novel, heretofore unseen Acrs.

**Figure 2.**
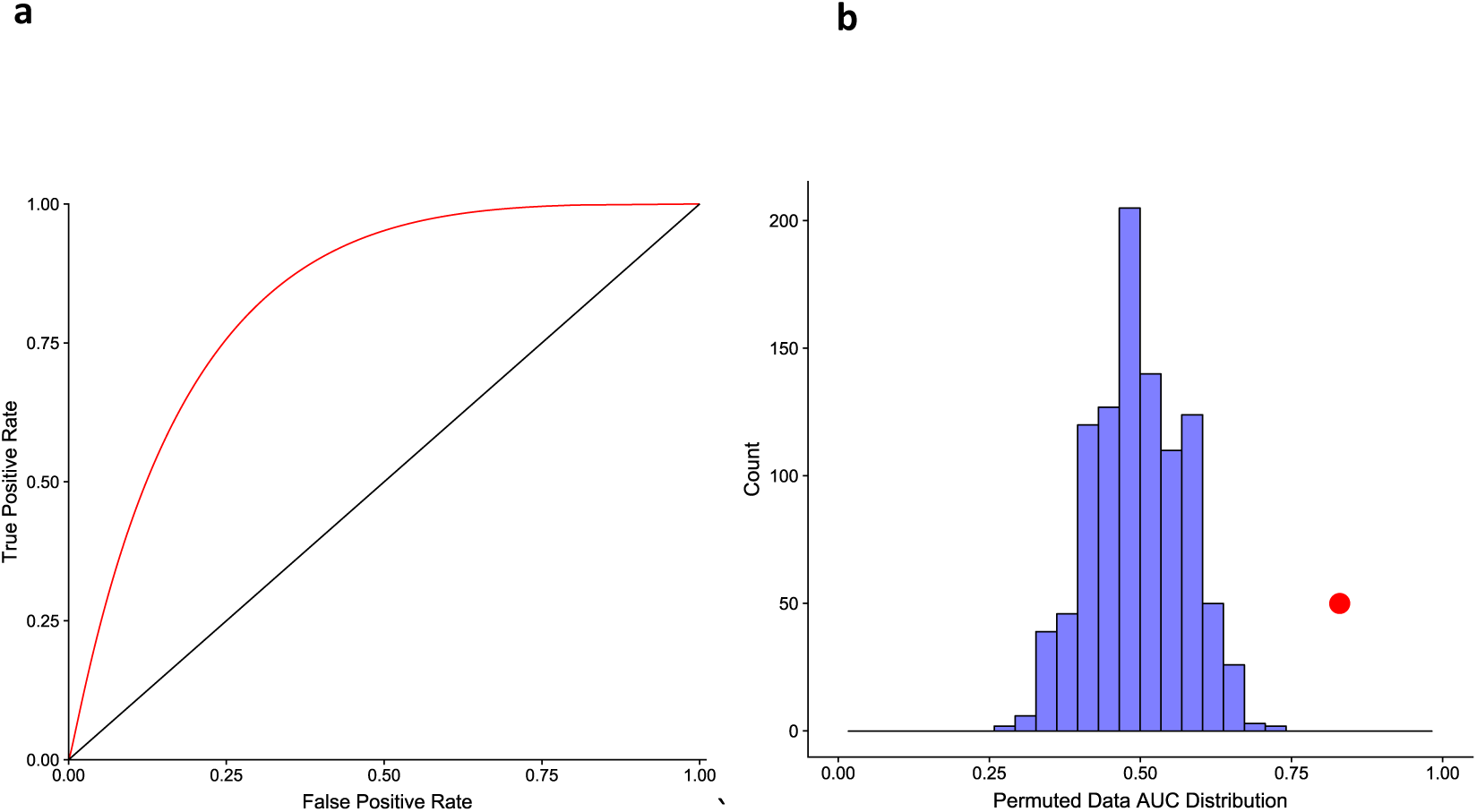
Model assessment on an unseen test set. A) The ROC AUC of the model scores on an unseen test set. B) A histogram of 1000 AUCs calculated using permuted model scores against the unseen test set representing the null AUC distribution. As expected for a well-calibrated assessment, the null AUC distribution is centered on 0.5, indicating random separation. The AUC for the correct model scores, 0.83, is indicated in red.

### Using the model to predict novel Acrs

Having demonstrated the predictive power of our model on the test set of recently discovered Acrs, we sought to predict novel Acrs. The first step in this direction was to define an appropriate search space of proteins likely enriched for Acrs. The initial dataset consisted of 182,561,570 proteins of which the majority (182,332,040) came from prokaryotes, and the rest were encoded by viruses (229,530).

Acrs are typically encoded either within prokaryotic virus genomes, or within prokaryotic genomic regions that appear to be integrated viruses (proviruses) or other mobile genetic elements (MGEs)^10, 19^. We therefore identified a subset of the prokaryotic database that consisted of genomes containing complete CRISPR-Cas systems^31^, under the premise that these genomes are more likely to encompass prophages with Acrs targeting the respective CRISPR-Cas variants^16, 20^. We further sought to limit the prokaryote protein set to proteins encoded by (predicted) proviruses. Although there are many methods for predicting complete proviruses and their boundaries, these fall short of comprehensive identification of provirus regions in prokaryotic genomes, primarily, because numerous proviruses are inactivated and partially deteriorated^32, 33^. Indeed, many of the known Acrs are encoded in the vicinity of virus proteins^12^ but not necessarily within clearly active proviruses encoding hallmark virus genes and bounded by well-defined provirus boundaries. Therefore, instead of explicitly predicting proviruses, we enriched for virus-related sequence, by filtering the prokaryote protein set to the proteins encoded in the vicinity of known virus proteins (see Methods for details). The resulting combined dataset of prokaryotic viruses and suspected proviruses consisted of 10,938,430 proteins. As these proteins are largely virus-related, we expected this set to be enriched for Acrs.

We assessed this set of proteins with our random forest model which resulted in an initial set of 1,546,505 candidate Acrs. We further filtered these to retain only those that had no significant hits to CDD^26^ or pVOG^27^, yielding 892,830 proteins (see Methods for details). This set of proteins was clustered by sequence similarity, resulting in 232,616 protein clusters (Supplementary File 2 available at ftp://ftp.ncbi.nih.gov/pub/wolf/_suppl/ACR20/supplementary_file_2.txt; see Methods for details). Heuristic filters were applied to each of these clusters, based on known Acr characteristics, to further enrich the candidate set for true Acrs (Table 2).

**Table 2.** Heuristics for filtering Acr families.

| Heuristic features of candidate protein clusters | Threshold |
| --- | --- |
| Number of members that have HTH-domain protein encoded downstream. | A minimum of one. |
| Number of members in self-targeting or virus genome. | A minimum of one. |
| Average directon size. | A maximum of 5 genes. |
| Number of homologs in our prokaryotic dataset. | A maximum of 374. |
| Ratio of prokaryotic homologs to suspected provirus homologs. | A maximum of 3 or a single virus homolog. |
| Number of HHBlits hits. | A maximum of 52. |

The hallmark characteristics of Acrs are that they i) are encoded upstream of HTH proteins, and ii) are found in self-targeting genomes^16^. We therefore required each family to have at least one member that fulfills each of these criteria. After this filtering, 11,304 families remained, among which 20 included known Acrs.

As genes encoding Acrs tend to form small directons, we sought to estimate a heuristic maximum threshold for the mean directon size in a candidate family that would enrich our protein set for true Acrs. We therefore searched for the threshold that, when applied, retained the largest fraction of the known Acrs in our set of 11,304 while filtering out as many of the candidate families as possible. To quantify this feature, we used the balanced accuracy metric, which is equal to the average of the fraction of correct classifications between the two groups. We found that a maximum mean directon size of 5 genes gave the highest balanced accuracy (see Methods for details). Consequently, we removed protein families with an average directon size of more than 5 genes. After this filtering, 5,507 families remained including 18 families of known Acrs.

To eliminate additional false positives, we PSI-BLASTed each protein family alignment against our sequence dataset and, under the premise that Acrs are highly variable, fast evolving proteins, removed families with numerous homologs in diverse prokaryotes. We found that the heuristic cutoff value for the number of prokaryote homologs that maximized balanced accuracy was 374. We therefore limited our set to clusters with no more than 374 significant hits to the prokaryotic protein set (Table 2). Next, we enriched for virus proteins by limiting to families that either include at least one homolog encoded in a virus genome or have a small ratio of prokaryote homologs to prophage homologs (see Methods for details). We found that the cutoff value for the prokaryote to provirus ratio that maximized balanced accuracy was 3 (Table 2). Finally, we sought to exclude families that have numerous annotations when assessed with HHBlits and thus include well-characterized non-Acrs^34^. We found the cutoff value that maximized balanced accuracy for the number of HHBlits hits was 52.

Although, by applying these heuristics, we likely filter out some true Acrs predicted by the model, we expect that, overall, this approach enriches the resulting protein set for true Acrs. After applying the above filters, our enriched set consisted of 2,526 protein families (Figure 3, Supplementary File 2).

**Figure 3.**
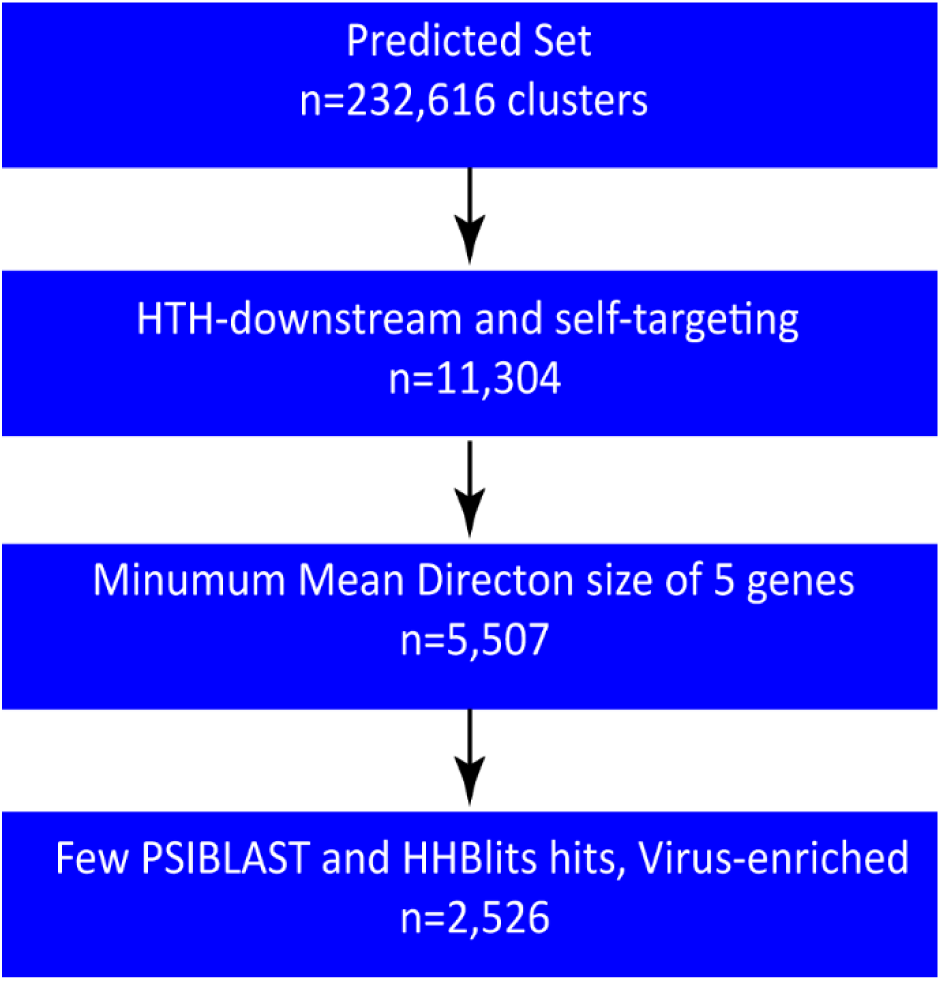
Flowchart illustrating the heuristic filtering.

### Characteristics of Predicted Acrs

We PSI-BLASTed all 2,526 candidate protein family alignments against a dataset of known Acrs and Acr-related sequences (see Methods for details). For 26 of these families, significant hits to the Acr set were detected. Of these protein families, 22 included known Acrs. The remaining 4 families with significant similarity to known Acrs are homologous to uncharacterized proteins (OrfA, OrfB, or OrfE) that are encoded within previously described Acr directons, namely, in the genomic neighborhoods of AcrIIA1-4 in *Listeria monocytogenes*, and all have been suspected of Acr activity although did not show Acr activity when tested^20^.

After removing these 26 families, we obtained 2,500 novel candidate Acr families, consisting of 16,919 putative Acrs. The mean size of a family was 7, the largest family included 319 members, and nearly half of the families (49%) were singletons (Figure 4).

**Figure 4.**
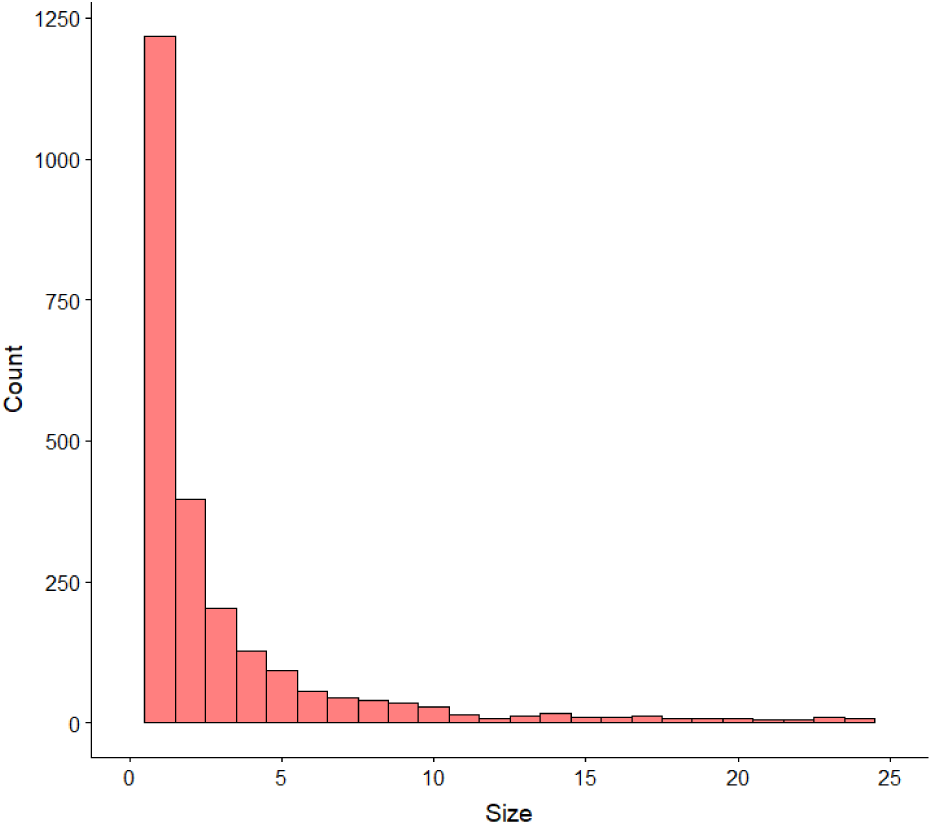
Histogram of predicted Acr family sizes. For visual clarity, families with more than 25 members are not displayed.

Given the different cluster sizes, each predicted Acr was assigned a weight inversely proportional to the size of its cluster, in order to ensure that related and highly similar predicted Acrs were not over-represented in summary statistics. Specifically, each predicted Acr was assigned a weight of 1/*n*, where *n* is the number of predicted Acrs in its cluster.

The predicted Acrs have a weighted average size of 109 aa, with a standard deviation (SD) of 71.6 (Figure 5A). As expected by design, the Acr genes tend to form small directons (weighted mean: 3.4; weighted SD: 1.47) consisting of short genes (weighted mean: 200aa; weighted SD: 155) (Figure 5B). The weighted mean isoelectric point of the predicted Acrs is 7.73 with a weighted SD of 2.6, and the weighted mean hydrophobicity is -0.31 with a weighted SD of 0.5. Per TMHMM and SignalP predictions^35, 36^, a weighted 15% of predicted Acrs have at least one putative transmembrane helix or signal peptide which, as expected, is substantially less than the expectation based on the negative set (28%, Table 1).

**Figure 5.**
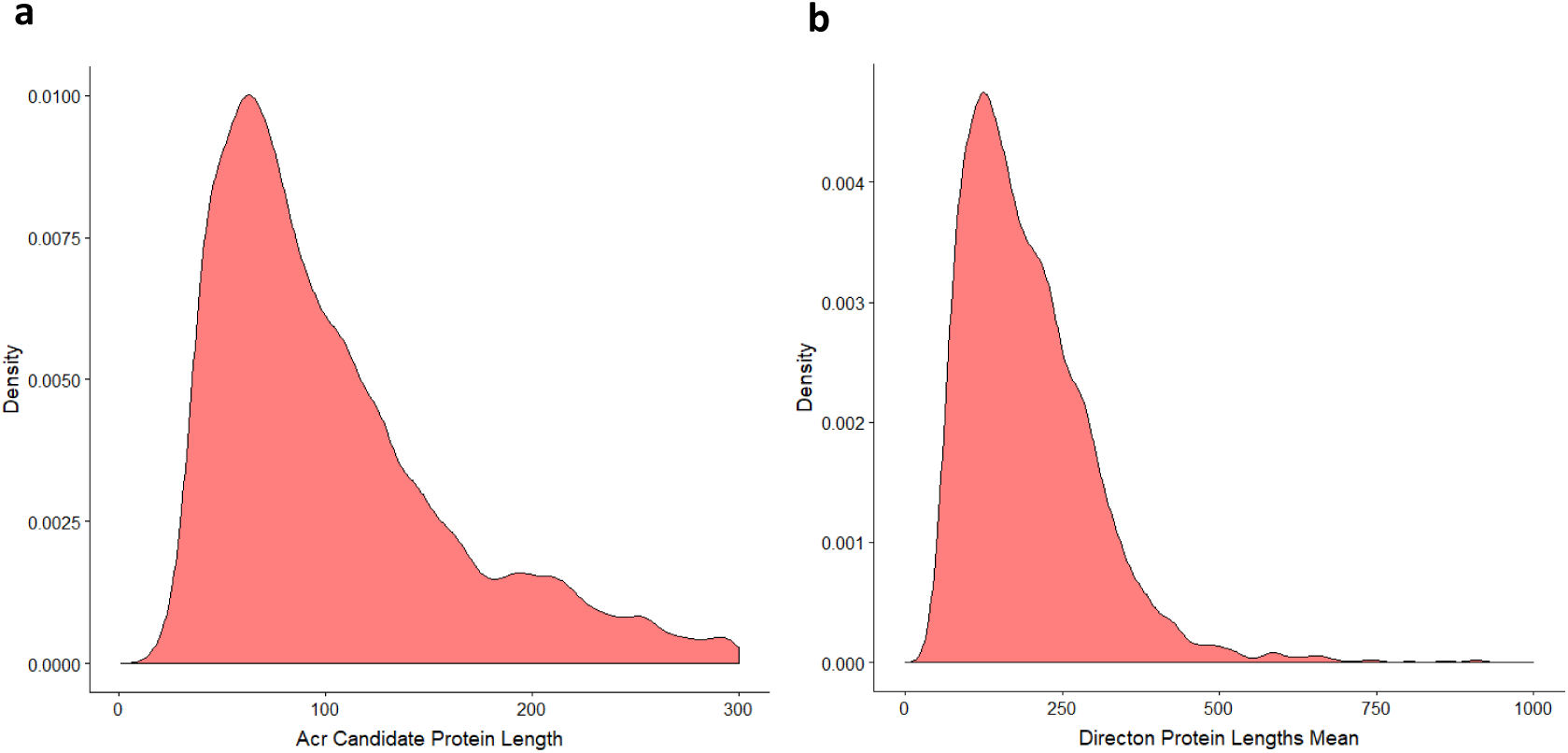
Protein length distribution of the Acr candidates. A) Density plot of the predicted Acrs protein lengths (mean: 109 aa) B) Density plot of the mean protein lengths of the predicted Acrs directons (mean: 200 aa).

Using JPred^37^, we predicted the secondary structure of the consensus sequences in the predicted Acr set. The mean percentage of amino acids contributing to alpha helices was 39%, and the mean percentage of amino acids contributing to beta sheets was 13%. These values did not differ significantly from the negative set, in which 97% and 85% of the proteins were predicted to contain at least one alpha helix or beta sheet, respectively, and the mean percentage of amino acids contributing to alpha helices and beta sheets was 41% and 13%, respectively.

The candidates are distributed across a diverse set of species (n=1,770). *Escherichia coli* accounts for the largest share of candidate Acrs at 2.37%. *Peptoclostridium difficile* (1.46%) and *Clostridium botulinum* (1.16%) round out the top three. Overall, each of the 1,770 species contains an average of 0.06% of the predicted Acrs. The most common CRISPR-Cas system in the genomes containing a predicted Acr(s) was subtype I-E (18.6%) followed by I-C (10.5%) and I-B (9.2%).

Among the candidate Acr clusters, 10% include at least one member encoded in a virus genome, with 279 virus strains encoding at least one Acr. Of the analyzed virus genomes, 197 (71%) encode a single predicted Acr, 66 (24%) encode two, and the remaining ones (5%) encode three or more Acrs. Archaeal viruses are also represented in this set, with 33 predicted Acrs predicted for 21 Archaeal viruses.

The maximum number of predicted Acrs in a single virus strain was 5, observed in Ruegeria phage DSS3-P1, four of which fell in the same HTH-containing directon. The viruses that were found to most commonly encode more than one Acr were *Mycobacterium* phages, followed by *Bacillus* and *Synechococcus* phages. Of the Archaeal viruses, the viruses that were found to most commonly encode more than one Acr were *Sulfolobales* Mexican rudivirus followed by *Sulfolobus islandicus* viruses.

We sought to examine the genomic context of the largest predicted Acr clusters and gauge how often they tend to appear in similar genomic neighborhoods. We examined the 10 largest Acr clusters and generated a presence-absence matrix for the members of these clusters in different genomic neighborhoods (Figure 6), with a genomic neighborhood defined as the 10 genes upstream and downstream of each Acr. Each column is a genomic neighborhood (ordered by similarity) and each row represents an Acr family (see Methods for details). Whereas the larger Acr clusters in this subset tend to appear in similar genomic neighborhoods, within these neighborhoods, we also find scattered predicted Acr singletons. This pattern is similar to what has been observed in known Acrs, where the Acrs present in a given directon vary across closely related strains, with some Acrs appearing in nearly all instances of the directon and others appearing sporadically^20^.

**Figure 6.**
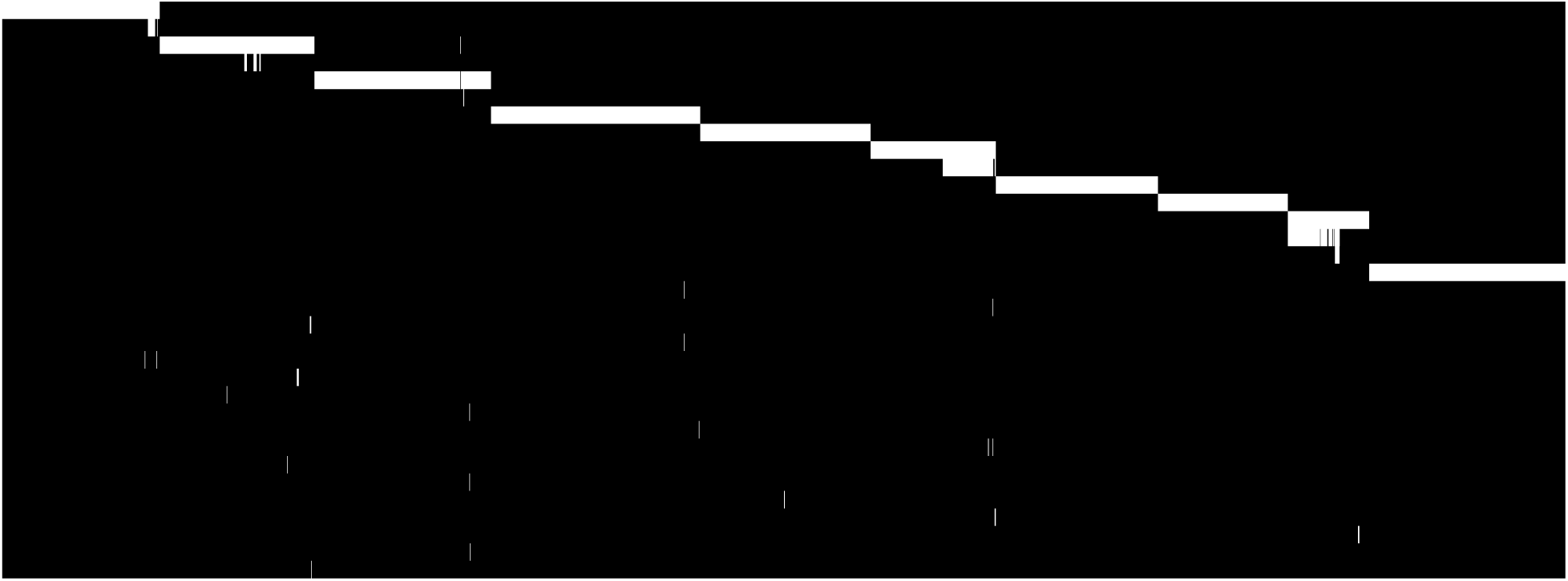
Presence=absence matrix of Acr families in genomic contexts. A binary matrix where each column is a distinct genomic neighborhood (ordered by similarity) and each row represents an Acr family (see Methods for details). Each cell represents the presence or absence of a member from the Acr family in the neighborhood, with grey representing presence and black representing absence.

### Case by case analysis of top Acr candidates

We next examined in greater detail the top candidates from our Acr candidate set. The top 30 families with at least 4 members were selected from the candidate set and explored using HHPRED^38^, PSI-BLAST against NR and examination of the genomic context for each candidate (see Methods for details). Supplementary Table 2 presents the key features of the top 30 candidates.

Here, we present in some detail the genomic contexts and characteristics of the top 5 candidates. It has been previously shown that Acrs tend to be encoded in short directons consisting of small genes, usually including one HTH gene^18, 20^. Further, they tend to fall in self-targeting genomes, usually in the vicinity of MGE proteins. This configuration has been seen with multiple Acr families across numerous species. One classic example of this is the Acr IIA1-4 families^20^ (Figure 7), which were shown to be encoded in such a configuration.

**Figure 7.**
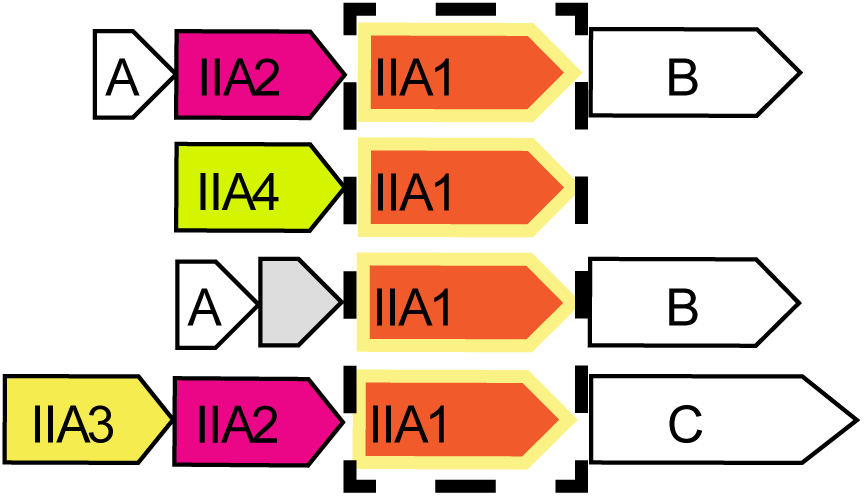
Genomic context of Acrs IIA1-4. The genomic contexts of Acrs IIA1-4 in *Listeria*, with Acrs encoded in short directons of small genes including one for a protein containing an HTH domain. Proteins containing an HTH domain are indicated with a yellow outline. Also shown are OrfA, OrfB and OrfC, proteins that have been suspected of Acr activity although did not show Acr activity when tested for Acr function^20^.

#### Candidate 4338

Members of one of our top 5 candidate Acr clusters, candidate 4338 (hereafter C4338) were found in suspected prophages and phages of *Listeria monocytogenes*, adjacent to AcrIIA1, with three quarters of the members of this family found in self-targeting genomes. At the time of our analysis, C4338 was not found to be homologous to any of the previously discovered AcrIIA genes. However, shortly after the completion of the analysis and while this manuscript was in preparation preliminary results on testing C4338 for an anti-CRISPR function have been reported independently^39^. C4338 has been identified as an anti-Cas9 protein (AcrIIA12), supporting the utility of our approach to discover novel Acrs.

#### Candidate 20391

Members of the C20391 cluster were identified in one phage (*Listeria* phage B054) and four suspected prophages (one in *Listeria innocua* and three in *Listeria monocytogenes*) (Figure 8A). All the prophage-encoded homologs were found in self-targeting genomes that carry CAS-II-A. Three of these genomes also carry CAS-I-B. All the prophage-encoded members of this cluster were found in bacterial genomes that also encoded AcrIIA1, and two of these also encoded Acrs IIA2 and IIA3. Given that all the genomes encoding proteins of this family encompass CAS-II-A, we predict that this is the target of its anti-CRISPR activity, although targeting of CAS-I-B is difficult to rule out.

**Figure 8.**
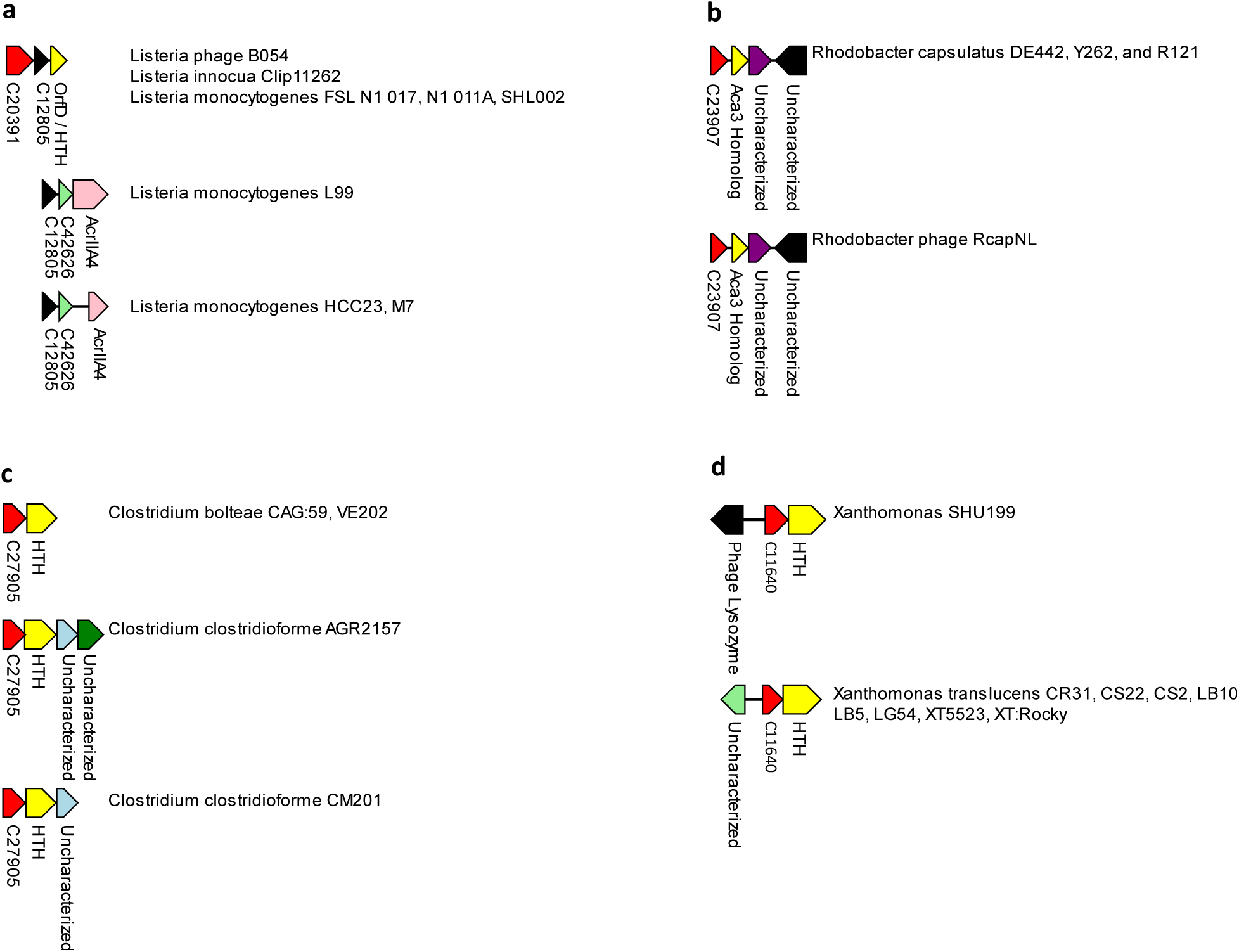
Genomic contexts of top candidates. A) Genomic context of C20391, C12805 and C42626 in 8 *Listeria* genomes. The genomic contexts of these three candidates are typical of the known Acrs, encoded in short directons of small proteins, with one protein including an HTH domain. B) Genomic context of C23907 in *Rhodobacter capsulatus* and *Rhodobacter* phage RcapNL. The genomic context of C23907 is typical of known Acrs and includes an HTH-containing protein that is a distant homolog of Aca3. C) Genomic context of C27905 in *Clostridium*. C27905 is found in directons between 2-4 proteins in length, and is adjacent to an HTH domain containing protein, as typical of known Acrs. D) Genomic context of C11640 in *Xanthomonas*. C11640 is found in directons consisting of 2 proteins, where the second protein is an HTH domain containing protein, as typical of known Acrs.

As is characteristic of known Acrs, C20391 homologs are typically encoded in short directons consisting of three genes (Figure 8A). One of these genes contains an HTH domain and is homologous to OrfD of *Listeria monocytogenes*. OrfD has been previously identified as a marker for Acr directons and is a distant homolog of AcrIIA1 although in itself, this protein has not been shown to possess Acr activity^20^. All members of this cluster are encoded adjacent to members of another predicted Acr family, C12805. C12805 includes 3 additional members that are not adjacent to C20391, but are all found in a directon with AcrIIA4 and an additional candidate, C42626, in prophages of *Listeria* strains solely containing CAS-I-B (Figure 8A).

In the genomic neighborhoods of C42626 (Figure 8A, *L. monocytogenes* L99, M7 and HCC23), one instance includes an expanded version of AcrIIA4 (encoded in *L. monocytogenes* L99) that contains an HTH, whereas the remaining two instances of AcrIIA4 lack an HTH domain. However, an examination of the nucleotide sequences immediately upstream of the truncated AcrIIA4 indicates that this truncation is likely to be an error in the sequence annotation, and that the N-terminal of these instances of AcrIIA4 can be extended to match the AcrIIA4 homolog in *L. monocytogenes* 99, including the HTH domain. Furthermore, the region of AcrIIA4 that contains the HTH domain is similar to the portion of OrfD that contains an HTH domain (38% identity), so that extended version of AcrIIA4 appears to be a fusion of OrfD and AcrIIA4.

Thus, candidates C20391, C12805, and C42626 all contain the hallmark characteristics of known Acrs, including their tendency to fall in known Acr neighborhoods and next to known Acr markers. This corroborating evidence greatly raises our confidence that these are true Acrs and further validates the predictive power of the methodology.

#### Candidate 23907

Members of the cluster C23907 were identified in one phage (*Rhodobacter* RcapNL) and in 3 RcapNL prophages integrated in self-targeting genomes of *Rhodobacter capsulatus*. C23907 belongs to a small directon of 3 genes, with the second gene in the directon containing an HTH domain. This HTH-containing gene is a distant homolog of Aca3, a previously discovered gene associated with Acrs, further supporting the prediction of anti-CRISPR functionality of C23907. The third gene in the directon is uncharacterized (Figure 8B). The 3 self-targeting prophages containing the Acr occur in genomes with two CRISPR systems, type I-C and type VI-A, either of which are potential targets of C23907.

#### Candidate 27905

Members of the C27905 cluster were found in *Clostridium*. Half of the homologs were found in genomes that are self-targeting. As is characteristic of Acrs, C27905 genes typically belong to a small directon of 2-4 genes, where the second protein encoded in this directon contains an HTH domain (Figure 8C). The other proteins in the directon are uncharacterized. All the genomes in this set contain CRISPR I-C, a potential target of C27905.

#### Candidate 11640

Members of the C11640 cluster were found in *Xanthomonas*. Eight of these genes were identified in *Xanthomonas translucens* and one in *Xanthomonas* sp. SHU199. Eight of the nine homologs were found in self-targeting genomes. C11640 tends to fall in a small directon of two genes, as is characteristic of known Acrs, where the second gene in the directon contains an HTH domain (Figure 8D). All the genomes containing C11640 have type I-C CRISPR systems, a potential target of C11640.

## Discussion

The Acrs are of major interest to a wide range of researchers, due both to their role in the evolutionary arms race between viruses and their prokaryotic hosts, and to their potential use as CRISPR-Cas inhibitors in genome engineering applications. However, identification of Acrs remains a formidable challenge, given their extreme variability and lack of functionally characterized homologs.

Here we demonstrate substantial predictive and discriminative power of a machine-learning approach for identification of candidate Acrs. This result appears unexpected given the paucity of distinctive features of the Acrs. Nevertheless, these few, rather generic features including the small size of the Acr genes, their arrangement in short directons that contain, additionally, genes for HTH proteins, poor evolutionary conservation, association with viruses and proviruses, and self-targeting seem to be sufficient for apparently robust Acr prediction. The underlying reason seems to be that, in viruses of prokaryotes, a substantial fraction, often, the majority of the genes that are not directly implicated in virus replication and morphogenesis are involved in anti-defense functions. A notable example can be found among archaeal viruses in some of which up to 40% of the genes appear to encode Acrs^40^. Hence a possible caveat of our predictions: some of the genes that we predict as Acrs might target other, non-CRISPR defense systems. Conversely, the possibility exists that, using the approach described here, we only detect one, albeit major, class of Acrs, whereas others might exhibit distinct properties. For example, a recently discovered Acr is an acetyltransferase that inhibits subtype V-A CRISPR-Cas via acetylation of the effector protein^13^. Another anti-V-A Acr is a nuclease that abrogates the CRISPR-Cas activity by cleaving the guide RNA^14^. Thus, a distinct class of enzymatically active Acrs seems to exist, and at least some of these can be larger and more evolutionarily conserved proteins than the Acrs addressed here.

The above caveats notwithstanding, the combination of sensitive database searches, machine learning and heuristic filters applied here yielded 2500 previously undetected families of strong Acr candidates that comprise an extensive resource, which we make accessible online (http://acrcatalog.pythonanywhere.com/), for structural and functional studies on Acr-CRISPR interactions, with likely subsequent applications. The current database of prokaryotic virus genomes is limited in scope but grows rapidly, thanks, largely, to metagenomic discovery of new viruses. Furthermore, so far, no targeted search for Acrs in MGEs other than viruses, such as plasmids or transposons, has been performed. Understanding the distribution of Acrs throughout the prokaryotic mobilome is a key next step to understanding the arms-race that can be expected to lead to the discovery of numerous novel Acrs. Thus, the clear next step is to extend our approach to search expanding virus genome databases, metagenomes, and other MGE. Iterative application of this strategy should greatly expand the diversity of Acrs.

## Methods

### Iterative Search for Acr Homologs

For each Acr family, we selected a single representative sequence and PSI-BLASTed it against NCBI’s non-redundant sequence database (NR). Iterative PSI-BLAST was run to convergence, the identified homologs were aligned using MUSCLE^41^, and the resulting alignment was PSI-BLASTed against our prokaryote dataset^24^ and our prokaryotic virus dataset from the NCBI viral genomes resource^42^. We used a cutoff of e-value less than or equal to 10e-4 for homolog detection and manually reviewed each resulting alignment.

### Weighting the Acrs

For the positive set, we sought to weight each Acr by its sequence similarity to the other Acrs, in order to avoid oversampling closely related data points. Initially, each Acr family is assigned a weight of one. Then, within each Acr family, its member proteins were clustered using mmseq2, with the parameters c=0.5 and s=0.4^43^. Each cluster is defined as a sub-family, and the initial weight of one given to the family is divided evenly amongst the sub-families. Following this, each sub-family’s weight is divided evenly among its members. Thus, each Acr’s weight is proportional to its similarity to other Acrs in the set.

For the negative set, an analogous procedure was followed. After randomly selecting a set of proteins as the negative set pool, these proteins were clustered using the same mmseq2 parameters as used for the Acr families, and from each cluster, a single representative was selected. Each representative protein was given a weight of one.

In training and in assessing the model, the negative set was re-weighted so that each class (Acr and non-Acr) had the same total weight.

### Protein Annotations

Proteins in our dataset were annotated by PSI-BLASTing protein alignments from CDD^26^ and pVOG^27^, with an e-value cutoff of 10e-4. Proteins with significant hits to to pVOGs were classified as viral.

### Self-Targeting Assemblies

Self-targeting assemblies were detected by BLASTing the spacers^31^ from each assembly in our dataset against the corresponding genome and filtering for exact matches. Wherever an exact match was found, the respective assembly was classified as self-targeting (Supplementary File 3).

### Defining The Features For The Model

Overall, 12 total features were defined. Some features related to the protein itself, while others relate to the protein’s directon. A directon was defined as consecutive proteins on the same strand with a maximum of 100bp between adjacent proteins.

The features were defined as follows:

*Protein size.* The length, in amino acids, of the candidate protein.

*Directon size.* The number of genes in the directon.

*Mean directon protein size.* The mean length, in amino acids, of all proteins in the directon.

*Protein hydrophobicity.* The protein’s hydrophobicity ^25^.

*Protein annotation.* A binary score of whether the protein is annotated or not. We consider a protein as “annotated” if it has at least one significant hit to any alignment, outside of alignments annotated as “hypothetical protein”, “putative predicted product”, or “provisional”.

*Fraction of directon that is annotated.* The fraction of proteins in the directon that are annotated as defined above.

*HTH-downstream.* Whether there is an HTH domain-containing protein encoded downstream of and adjacent to (within three genes) the Acr candidate within the same directon. This feature was analyzed by PSI-BLASTing proteins against the subset of alignments from the PVOG and CDD datasets containing in their name or description either the term “HTH” or “helix-turn-helix”, with an e-value cutoff of 5e-3.

*Self-targeting.* Whether the protein is encoded in a self-targeting genome.

*Predicted membrane association.* Whether the gene is predicted to be transmembrane or contain a signal peptide using tmhmm and signalp, respectively ^35, 36^.

*Fraction of membrane-associated proteins in directon.* The fraction of the proteins encoded in the directon that are predicted to be transmembrane or contain a signal peptide as defined above.

*Directon spacing.* The mean spacing between genes in the directon.

*Whether genome is viral.* Whether the protein is encoded in a viral genome or in a prokaryotic genome.

Ten generations of genetic feature selection were run, yielding the following best feature set:

1. Containing Genome is Self-Targeting
2. Directon Annotated Protein Fraction
3. Directon Protein Lengths Mean
4. Directon Size
5. Protein is Annotated
6. Protein has HTH-Downstream
7. Protein Length
8. Protein Hydrophobicity

### Building the Model

The model was constructed using scikit-learn (https://scikit-learn.org), specifically, the ExtraTreesClassifier with the the *n_estimator* parameter set to 1000, meaning that the random forest consisted of 1000 trees. The rest of the parameters were left at default.

The model was trained on the training data set described above, while down-weighting the negative set so that each class (Acr and non-Acr) has the same total weight.

Predictive scores were calculated by using the ExtraTreesClassifier function *predict_proba*. When calculating binary predictions, the threshold was set to the best value for differentiation in the training set when maximizing accuracy, which was equal to 0.09.

### Defining the Acr Search Space

#### Predicted Provirus Sequences

The alignments of the pVOG proteins were compared to the dataset of genomes containing CRISPR-Cas^24, 31^. Each directon containing a protein with a viral hit with an e-value below 10e-4 was considered a provirus-related sequence, along with the adjacent directons on either side. Adjacent blocks of prophage-related directons (within 500 bp of each other) were considered as provirus candidates. If the provirus candidate contained at least two virus hits within 3 kb of each other, it was considered a predicted prophage.

#### Prokaryotic Virus Sequences

The set of virus proteins was assembled from the NCBI viral genomes resource^42^ and subset to prokaryotic viruses based on taxonomy data (https://www.ncbi.nlm.nih.gov/genomes/GenomesGroup.cgi?taxid=10239). This virus set totaled 229,530 proteins encoded in 2,291 genomes.

### Permutation P-Value Calculation

To calculate permutation p-values, the model’s predictions for the test set were shuffled. We then tested how well the model performed on this shuffled dataset. This procedure was repeated 1000 times, creating a null distribution of AUCs. With this null distribution, a permutation p-value was calculated as follows. Let *n_p_* be the number of AUCs in the null distribution that are greater than or equal to the actual observed AUC. The permutation p-value, then, is equal to 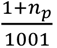. Thus, when the actual AUC was greater than any AUC in the entire permuted set, the p-value was approximately 0.001.

### Clustering and Weighting Candidate Acrs

Candidate Acrs were clustered using mmseq2, with the parameters c=0.5 and s=0.4^43^. A weight of 1/nc was assigned to each cluster, where n_c_ is the number of Acr candidate clusters. The weight of each cluster was then divided evenly among all protein members of the cluster, so that the weight of each Acr was inversely proportional to the size of the cluster it belonged to. These weights were used when calculating summary statistics for the Acr candidate set, to avoid oversampling closely related data points.

### PSI-BLASTing Against Known Acr and Acr-related Sequences

We created a sequence database of known Acrs and Acr-related sequences (Supplementary File 4). This database included all known Acrs, Acas, and proteins previously suspected of possessing Acr activity but not showing any when tested. We included the group of previously suspected Acr proteins as these are proteins that bear Acr characteristics, and therefore may be detected by our method, but have already been tested for Acr activity.

The alignment of each candidate Acr cluster was PSI-BLASTed against this dataset of known sequences, the clusters that produced hits with an e-value of less than 10e-3 were discarded as belonging to known Acr families or families that have already been already tested for the Acr function.

### Heuristic Filtering

To choose the thresholds for all the heuristics except for self-targeting and HTH-downstream, 10 evenly spaced threshold values were tested, between the minimum Acr value and the maximum Acr value.

Each of these 10 thresholds were applied as cutoffs to the Acr families, and for each threshold the balanced accuracy was calculated. The balanced accuracy is equal to the mean of the percentage of known Acrs that passed the threshold and the percentage of all proteins that were filtered by the threshold, so that a higher balanced accuracy corresponds to better discrimination between the known Acrs and the rest of the candidates. The final threshold was selected so as to maximize the balanced accuracy. The selected threshold was then applied to the dataset.

Six heuristics were defined to further enrich the Acr candidate set.

*Number of members that have HTH-downstream.* We required that at least one member of the candidate family have an HTH-containing protein encoded downstream within the same directon.

*Number of members in self-targeting or virus genome.* We required that at least one member of the candidate family was either encoded in a self-targeting genome or encoded in a virus genome.

*Mean directon length.* The mean number of genes in the directon for all members of the family.

*Number of homologs in prokaryotic dataset.* The multiple protein alignment of each family was PSI-BLASTed against the prokaryotic sequence dataset^24^ and filtered for hits with a maximum e-value of 10e-6, 50% identity and 50% query coverage. All families with more than 400 hits were discarded.

*Ratio of prokaryotic homologs to predicted provirus homologs.* Multiple protein alignment of each family was PSI-BLASTed against the predicted provirus sequence dataset and the virus sequence dataset and filtered for hits with a maximum e-value of 10e-6, 50% identity and 50% query coverage.

If a family produced at least one hit to a virus sequence, it was included. If not, it was required that the ratio between the number of hits to the prokaryotic sequence dataset to the number of hits to the predicted provirus dataset was less than or equal to three.

*Number of HHBlits hits.* The alignment of each family was compared to PFAM^44^ and PDB70^45^ using HHBlits^34^. Families with more than 100 hits were discarded.

### Construction of Acr Presence-Absence Matrix

To generate the presence-absence table, for the 10 largest Acr clusters, 10 genes upstream and downstream were extracted where available (a maximum of 20 genes total). If within this set, an additional predicted Acr was represented, the set was further extended to include the 10 genes upstream and downstream of that additional predicted Acr. The resulting gene arrays were considered the Acr genomic neighborhood.

A binary matrix was constructed where each column is a genomic neighborhood, ordered by content similarity, and each row is a predicted Acr family. In addition to the Acrs from the top 10 largest clusters, those encoded within 10 genes upstream or downstream of Acrs from the largest clusters were included. Each cell represents the presence or absence of a member of the respective Acr family in the neighborhood.

### Manual Assessment of Candidates

The multiple alignment for each of the top 30 candidates in Supplementary Table 2 was compared against the PDB, PFAM and NCBI CD databases using HHPRED^38^. For each candidate we calculated a consensus sequence, where the consensus letter for an alignment position was defined as the amino acid that has the highest BLOSUM62 score among the amino acids occupying the position. The consensus sequence of each candidate family was PSI-BLASTed against NR, and the genomic contexts of homologs were visually assessed using Geneious Prime.

## Acknowledgements

The authors thank Koonin group members for helpful discussions. This research was supported by the Intramural Research Program of the National Library of Medicine at the NIH.

## Author Contributions

A.B.G., K.S.M., Y.I.W., and E.V.K. designed research; A.B.G. performed research; A.B.G., S.A.S., K.S.M., Y.I.W., J.B.-D., and E.V.K. analyzed data; and A.B.G. and E.V.K. wrote the paper.

## Competing Interests

J.B.-D. is a scientific advisory board member of SNIPR Biome and Excision Biotherapeutics and a scientific advisory board member and co-founder of Acrigen Biosciences.

